# Sensorimotor feedback loops are selectively sensitive to reward

**DOI:** 10.1101/2021.09.16.460659

**Authors:** Olivier Codol, Mehrdad Kashefi, Christopher J. Forgaard, Joseph M. Galea, J. Andrew Pruszynski, Paul L. Gribble

**Affiliations:** Brain and Mind Institute, University of Western Ontario, Ontario, Canada; Department of Psychology, University of Western Ontario, Ontario, Canada; School of Psychology, University of Birmingham, United Kingdom; Department of Physiology & Pharmacology, Schulich School of Medicine & Dentistry, University of Western Ontario, Ontario, Canada; Robarts Research Institute, University of Western Ontario, Ontario, Canada; Haskins Laboratories, New Haven CT, USA

## Abstract

Although it is well established that motivational factors such as earning more money for performing well improve motor performance, how the motor system implements this improvement remains unclear. For instance, feedback-based control, which uses sensory feedback from the body to correct for errors in movement, improves with greater reward. But feedback control encompasses many feedback loops with diverse characteristics such as the brain regions involved and their response time. Which specific loops drive these performance improvements with reward is unknown, even though their diversity makes it unlikely that they are contributing uniformly. We systematically tested the effect of reward on the latency (how long for a corrective response to arise?) and gain (how large is the corrective response?) of seven distinct sensorimotor feedback loops in humans. Only the fastest feedback loops were insensitive to reward, and the earliest reward-driven changes were consistently an increase in feedback gains, not a reduction in latency. Rather, a reduction of response latencies only tended to occur in slower feedback loops. These observations were similar across sensory modalities (vision and proprioception). Our results may have implications regarding feedback control performance in athletic coaching. For instance, coaching methodologies that rely on reinforcement or “reward shaping” may need to specifically target aspects of movement that rely on reward-sensitive feedback responses.

## 1. Introduction

If a cat pushes your hand whilst you are pouring a glass of water, a corrective response will occur that acts to minimise spillage. This simple action is an example of a behavioural response triggered by sensing a relevant change in the environment—here, a push that perturbs the movement of your arm away from the intended movement goal. This form of feedback control requires the brain to integrate sensory information from the periphery of the body, and thus suffers from transmission delays inherent to the nervous system. However, there is evidence that when more is at stake, we react faster to respond to demands of the task (Reddi and Carpenter, 2000). For instance, if wine was being poured instead of water, and your favourite tablecloth covers the table, you may be faster at correcting for a perturbation that risks spilling your wine.

In the context of human motor control, feedback-based control is not a monolithic process (Reschechtko and Pruszynski, 2020; Scott, 2016). Rather, the term encompasses a series of sensorimotor feedback loops that rely on different sensory information, are constrained by different transmission delays (Figure 1), and are supported by different neural substrates (Reschechtko and Pruszynski, 2020). For instance, the circuitry underlying the short-latency rapid response is entirely contained in the spinal cord (Liddell and Sherrington, 1924). The long-latency rapid response relies on supraspinal regions such as the primary motor and primary sensory cortices (Cheney and Fetz, 1984; Day et al., 1991; Evarts and Tanji, 1976; Palmer and Ashby, 1992; Pruszynski et al., 2011a), and is modulated by upstream associative cortical regions (Beckley et al., 1991; de Graaf et al., 2009; Omrani et al., 2016; Takei et al., 2021; Zonnino et al., 2021). Visuomotor feedback responses rely on visual cortex and other cortical and subcortical brain regions (Day and Brown, 2001; Desmurget et al., 2004; Pruszynski et al., 2010). Due to these differences, each feedback loop is governed by different objectives such as maintenance of a limb position or reaching toward a goal (Figure 1). Therefore, to address whether sensorimotor feedback is sensitive to motivational factors requires testing multiple distinct feedback responses. In this work, we employed rewarding outcomes (specifically, monetary reward) as a means to manipulate motivation (Codol et al., 2020a; Galea et al., 2015; Goodman et al., 2014; Hübner and Schlösser, 2010; McDougle et al., 2021).

**Figure 1:**
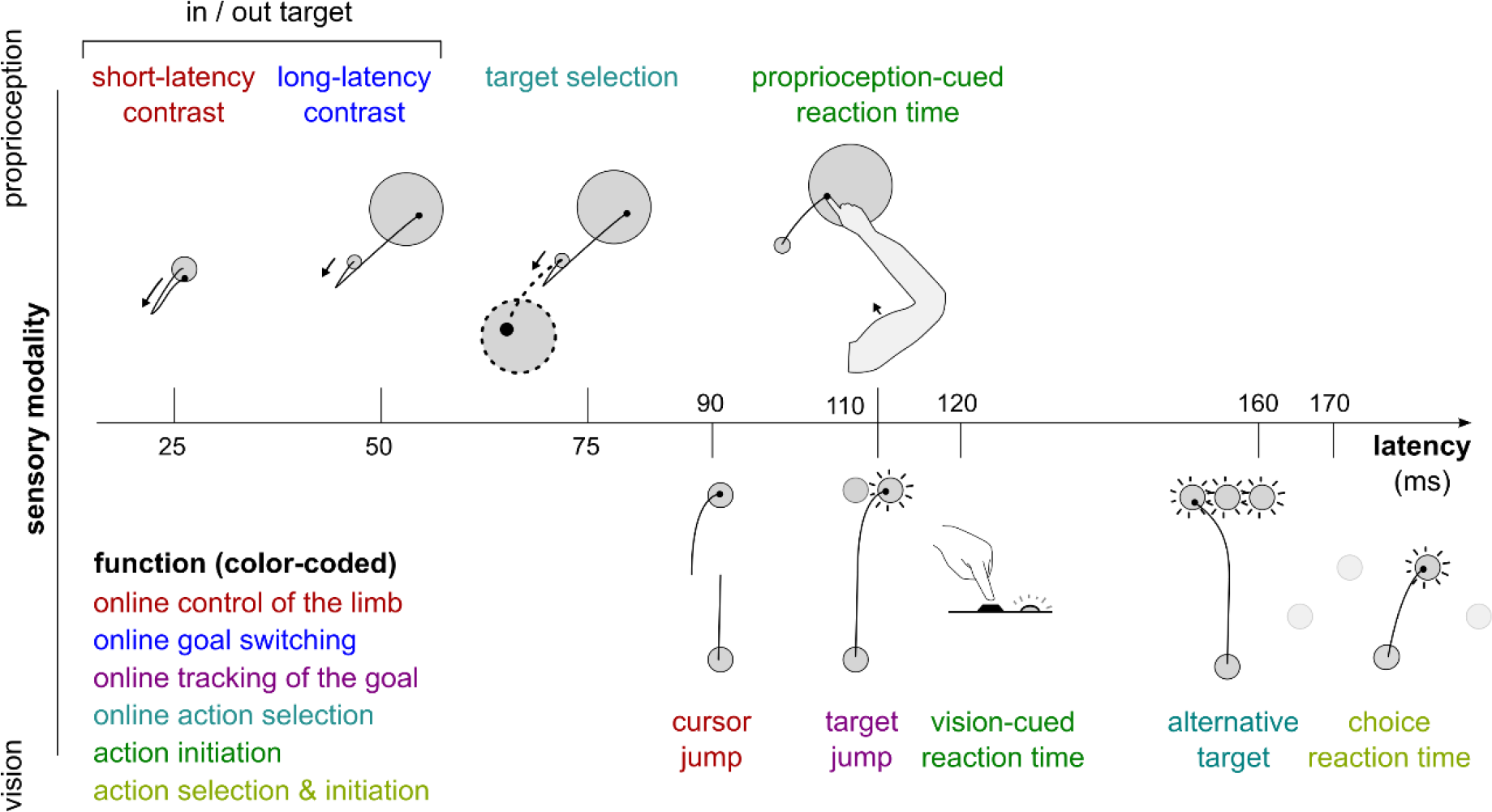
Different sensorimotor feedback responses are emphasized in different task designs. Feedback responses can be classified along three dimensions: the sensory modality on which they rely (vertical axis), their post-perturbation latency (horizontal axis), and the function they perform (color-coded). Latencies indicated here reflect the fastest reported values from the literature and not necessarily what was observed in this study. Note that this is a partial inventory. Figure adapted from Scott (2016).

Recent work has demonstrated that rewarding outcomes improve motor performance in many ways. Reward results in changes to the speed-accuracy trade-off, a hallmark of skilled performance (Codol et al., 2020a, 2020b; Manohar et al., 2015, 2019). Reward can lead to a reduction in noise in the central nervous system and at the effector to improve the control of movement (Codol et al., 2020a; Goard and Dan, 2009; Manohar et al., 2015; Pinto et al., 2013). But whether reward modulates sensorimotor feedback control specifically remains scarcely tested, although previous work in saccadic eye movements (Manohar et al., 2019) and indirect evidence in reaching (Codol et al., 2020a) suggests this may be the case. More recent studies outline a general sensitivity of feedback control to reward during reaching but does not differentiate between each distinct feedback loop that the nervous system relies on to implement this control (Carroll et al., 2019; De Comité et al., 2021; Poscente et al., 2021). However, the information to which each loop is tuned greatly varies (Reschechtko and Pruszynski, 2020; Scott, 2016), and consequently it is unlikely that they are all uniformly impacted by reward.

Systematically assessing each individual feedback loop’s sensitivity to reward provides a mechanistic, rather than descriptive, understanding of this relationship. Additionally, it further completes the picture of how rewarding information improves motor performance, a point of focus in the past decade due to applications in clinical rehabilitation (Goodman et al., 2014; Quattrocchi et al., 2017), sports coaching (Cashaback et al., 2019; Galea et al., 2015; Manley et al., 2014; Parvin et al., 2018), or studies on motor pathologies (Izawa and Shadmehr, 2011; Manohar et al., 2015, 2019; Pekny et al., 2015; Therrien et al., 2016, 2018). Teasing apart which feedback loops are modulated by reward can also provide a window into the neural substrates of reward processing during the production of movement, since different feedback loops are mediated by different brain regions (Omrani et al., 2016; Reschechtko and Pruszynski, 2020; Takei et al., 2021).

In the present study, we tested how seven distinct sensorimotor feedback responses are modulated by reward. We measured feedback latency (how long does it take for a corrective response to arise) and feedback gain (how large is the corrective response) for each feedback response within rewarded and unrewarded conditions. Motivational factors can take different forms, such as rewarding or punishing outcomes (Chen et al., 2017, 2018a, 2018b; Codol et al., 2020a; Galea et al., 2015; Guitart-Masip et al., 2014), inhibition versus movement (Chen et al., 2018a; Guitart-Masip et al., 2014), contingency (Manohar et al., 2017), expectation (Lowet et al., 2020; Schultz et al., 1997), urgency (Poscente et al., 2021), or agency (Parvin et al., 2018). In this study, we focused on contingent rewarding outcomes, in which participants have agency over the returns they obtain, and with an expectation component since potential for returns is indicated at the start of each trial (see Results and Methods).

## 2. Results

We first assessed feedback gain and latency for the short- and long-latency rapid response (SLR and LLR), which are the fastest feedback responses observed in human limb motor control. Participants were seated in front of a robotic device that supported their arm against gravity and allowed for movement in a horizontal plane. They positioned their index fingertip at a starting position while countering a +2 Nm background load (dashed arrows in Figure 2a, middle panel) to activate the elbow and shoulder extensor muscles. We recorded electromyographic signals (EMG) using surface electrodes placed over the brachioradialis, triceps lateralis, pectoralis major (clavicular head), posterior deltoid, and biceps brachii (short head). After participants held their hand in the starting position for 150-200 ms, a 10 cm radius target appeared at 20 degrees either inward (closer to the chest) or outward (away from the chest) with respect to the elbow joint. Next, a ±2 Nm torque perturbation was generated by the robot about the elbow and shoulder joints (solid arrows in Figure 2a, middle and right panels). A positive or negative torque signifies an inward or an outward perturbation from the starting position, respectively. Participants were instructed to move their fingertip into the target as soon as possible after the perturbation occurred. That is, the perturbation acted as a cue to quickly move the hand into the displayed target. This yielded a 2×2 factorial design, in which an inward or outward perturbation was associated with an inward or outward target (Pruszynski et al., 2008). In the present study we will refer to this task as the “In-Out Target” task. Different contrasts allowed us to assess the SLR and LLR within this task (Figures 2a and 3a).

**Figure 2:**
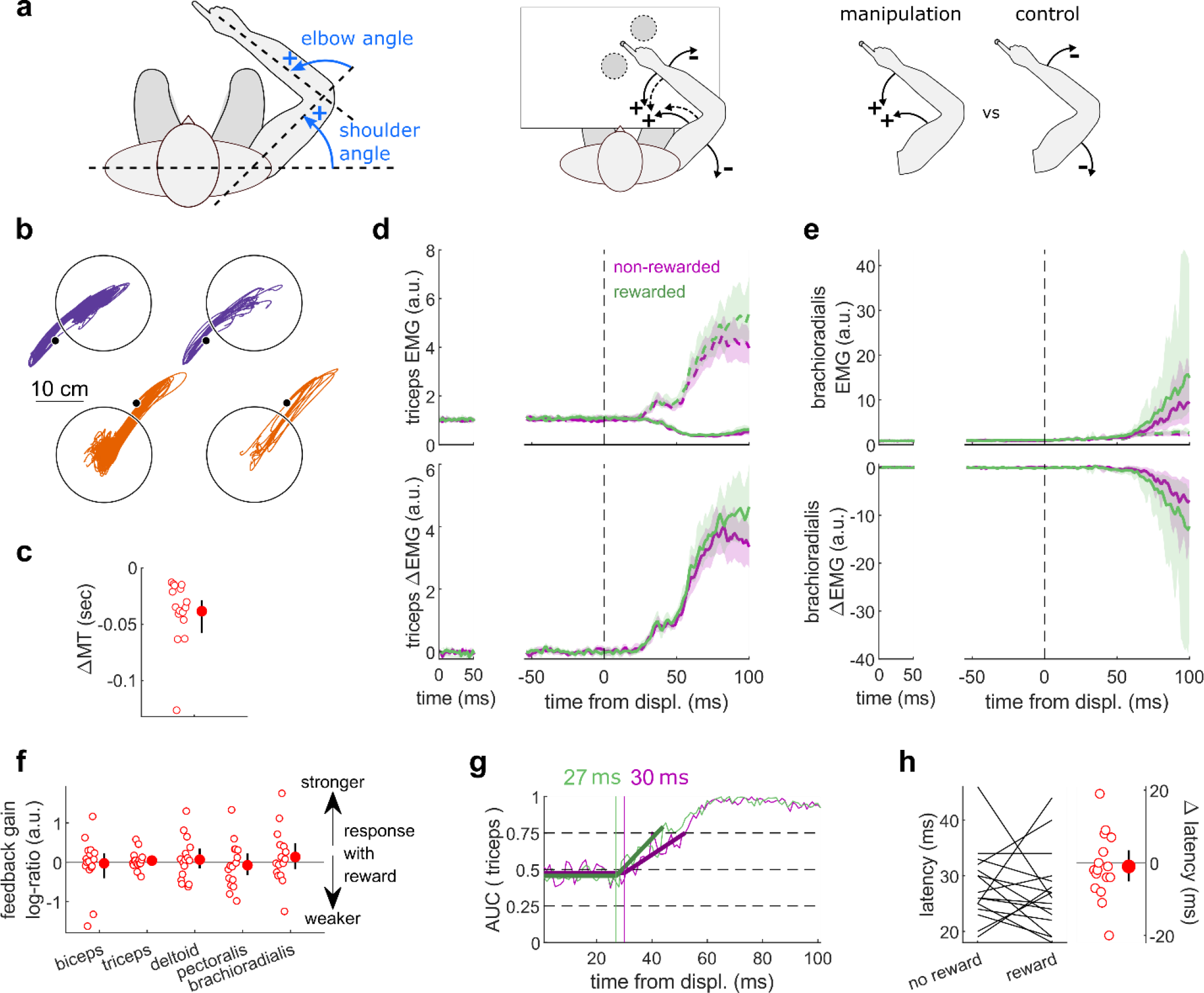
Results for the SLR contrast. **(a)** Left panel: representation of the joint angle definition employed in this study. Middle panel: top view schematic of the apparatus. Participants could move their arm in a horizontal plane. Background forces were applied to pre-activate the extensor muscles (dashed arrows) and a mechanical perturbation, either two positive or negative torques, was applied to elicit the SLR and LLR (solid arrows). The dashed circles indicate the two possible target positions. Right panel: contrast used to observe the SLR. Background loads are not drawn here for clarity. **(b)** Trajectories for one participant (left) and average trajectory for all participants (right, N=16). **(c)** Difference in mean movement time between rewarded and non-rewarded trials. A negative value indicates a smaller movement time for rewarded trials. **(d)** Average triceps EMG signal across participants, with the dashed and solid lines representing manipulation and control conditions, respectively, as indicated in (a); bottom panels: difference between the manipulation and control condition. The left panels show EMG at trial baseline (see Methods). Shaded areas indicate 95% CIs. **(e)** Same as (d) but for the brachioradialis. **(f)** Feedback gains following SLR onset. **(g)** Example area under curve (AUC) to obtain response latency for one participant. Thick lines indicate line-of-best-fit for a two-step regression (see methods). **(h)** Response latencies. In all panels with a red filled dot and black error bars, the filled dot indicates the group mean and error bars indicate 95% CIs.

**Figure 3:**
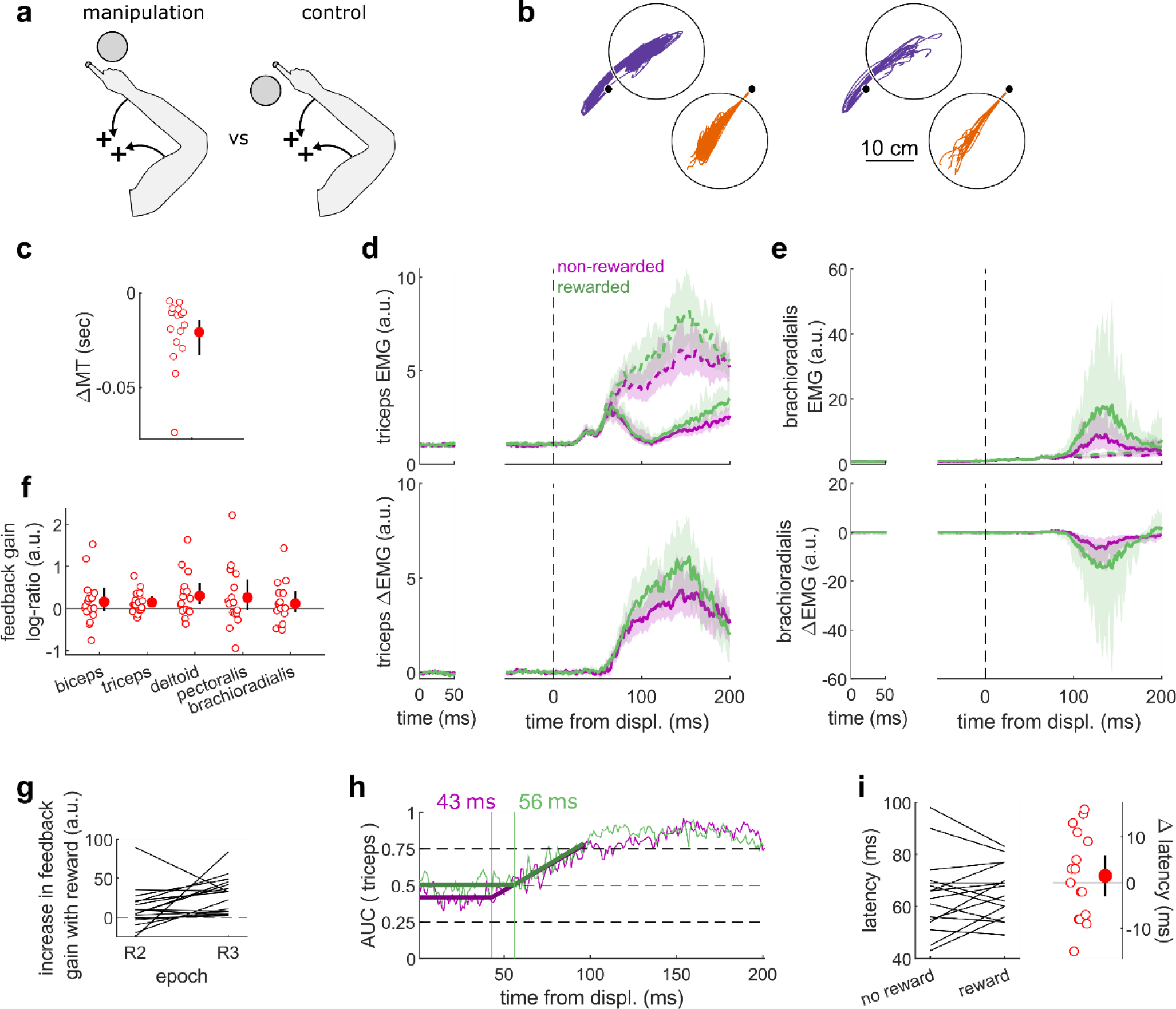
Results for the LLR contrast. **(a)** Contrast used to observe the LLR. Background loads are not drawn here for clarity. **(b)** Trajectories for one participant (left) and average trajectory for all participants (right, N=16). **(c)** Difference in mean movement time between rewarded and non-rewarded trials. A negative value indicates a smaller movement time for rewarded trials. **(d)** Average triceps EMG signal across participants, with the dashed and solid lines representing manipulation and control conditions, respectively, as indicated in (a); bottom panels: difference between the manipulation and control condition. The left panels show EMG at trial baseline (see methods). Shaded areas indicate 95% CIs. **(e)** Same as (d) but for the brachioradialis. **(f)** Feedback gains following LLR onset. **(g)** Difference in triceps LLR feedback gain between reward and no-reward conditions for each participant, in the R2 and R3 epochs. **(h)** Example area under curve (AUC) to obtain response latency for one participant. Thick lines indicate line-of-best-fit for a two-step regression (see methods). **(i)** Response latencies. In all panels with a red filled dot and black error bars, the filled dot indicates the group mean and error bars indicate 95% CIs.

Unlike previous studies using this paradigm, we introduced monetary reward as a third factor to assess its impact on feedback responses. Rewarded and non-rewarded trials were indicated at the beginning of each trial by displaying “$$$” and “000” symbols, respectively, on the display screen in front of participants. These symbols were replaced by each trial’s actual monetary value once the target was reached (always 0 ¢ in the case of non-rewarded trials). For rewarded trials, the monetary gains were proportional to the time spent inside the end target, therefore promoting faster reaches (see Methods) because trial duration was fixed.

Feedback latency and gain were assessed by measuring when the EMG signal diverged between those two conditions, and the magnitude of the EMG signal following this divergence, respectively.

### 2.1. The short latency rapid response remained similar in rewarded and non-rewarded conditions

The SLR can be assessed by contrasting the trials with an inward perturbation versus those with an outward perturbation (Figure 2a, right panel). Figure 2b shows trials falling in these categories for a single participant (left hand side) as well as each participant’s average trajectory (N=16; right hand side). Note that the trials for which the target is in the direction of the perturbation were excluded from this visualisation for clarity, but in practice they are included in the analyses and their inclusion or exclusion does not have a significant impact on the results. Before comparing the impact of rewarding context on feedback responses, we tested whether behavioural performance improved with reward by comparing movement times (MT) expressed. To do so, we computed for each participant the median MT of trials corresponding to the conditions of interest (Figure 2a, rightmost panel) for rewarded and non-rewarded trials and compared them using a Wilcoxon rank-sum test. Indeed, median MTs were faster during rewarded trials than in non-rewarded ones (W=136, r=1, p=4.38e-4, Figure 2c).

The contrast successfully emphasizes the divergence in EMG response at the triceps (*triceps lateralis*, Figure 2d). We also displayed the antagonist brachioradialis muscle for comparison (Figure 2e). This allowed us to quantify feedback gains as a log-ratio of integrals of EMG activity between rewarded and non-rewarded conditions (see Methods), meaning a positive number indicates an increase in feedback gain for the rewarded conditions (Figure 2f). For all muscles, we observed no difference in a 25 ms window after the onset of the SLR (biceps: W=78, r=0.57, p=0.61; triceps: W=81, r=0.6, p=0.5; deltoid: W=76, r=0.56, p=0.68; pectoralis: W=83, r=0.61, p=0.44; brachioradialis: W=80, r=0.59, p=0.53, Figure 2f). Next, we assessed the time of divergence of each participants’ EMG activity between the reward and no-reward conditions using a Receiver Operating Characteristic (ROC) signal discrimination method (Figure 2g; Weiler et al., 2015). We performed this analysis on the rewarded and non-rewarded trials separately, yielding two latencies per participant. Latencies for the SLR in the non-rewarded conditions were in the 25 ms range post-perturbation, and latencies in the rewarded conditions were in a similar range as well, with no significant difference observed (W=70.5, r=0.52, p=0.57, Figure 2h). Therefore, rewarding outcomes affected neither feedback latency nor feedback gains of the SLR.

### 2.2. Reward altered feedback responses as early as 50 ms post-perturbation

The LLR typically arises more strongly if the direction of a mechanical perturbation to the limb conflicts with the task goal (the target; Pruszynski et al., 2008). Therefore, turning to the LLR, we contrasted trials with an inward perturbation and an outward target with trials with an inward perturbation as well but an inward target instead (Figure 3a-b). We performed that contrast for non-rewarded trials, and then for rewarded trials independently. As a control, we compared each participant’s median MTs in all the above trials (Figure 3a) with a rewarding context to those with a non-rewarding context. We observed that MTs were shorter in rewarded trials (W=131, r=0.96, p=1.1e-3, Figure 3c).

Feedback gains were greater in the rewarded condition for the triceps, deltoid, and brachioradialis in a 50 ms window following LLR onset (biceps: W=98, r=0.72, p=0.12; triceps: W=136, r=1, p=4.4e-4; deltoid: W=135, r=0.99, p=5.3e-4; pectoralis: W=96, r=0.71, p=0.15; brachioradialis: W=129, r=0.95, p=1.6e-3; Figure 3d-f). This 50 ms epoch is often split into an R2 and R3 epoch corresponding to its first and last 25 ms, because the processes contributing to the response in those two epochs tend to differ, as well as their neural substrates (Lee and Tatton, 1975; Pruszynski et al., 2011b; Tatton et al., 1975). We did observe an increase in feedback gains for the rewarded condition in the R2 epoch for the triceps and deltoid muscles (biceps: W=89, r=0.65, p=0.28; triceps: W=108, r=0.79, p=0.039; deltoid: W=108, r=0.79, p=0.039; pectoralis: W=93, r=0.68, p=0.2; brachioradialis: W=81, r=0.6, p=0.5). The R3 epoch showed an increase in feedback gains as well (biceps: W=100, r=0.74, p=0.098; triceps: W=136, r=1, p=4.4e-4; deltoid: W=131, r=0.96, p=1.1e-3; pectoralis: W=95, r=0.7, p=0.16; brachioradialis: W=129, r=0.95, p=1.6e-3). To assess whether this effect of reward is different in the R2 and R3 epoch, we tested for an interaction between reward and epoch by comparing the difference between triceps EMG signals in those epochs in the rewarded and unrewarded conditions. We focus on the triceps because the task emphasizes triceps activity and so yields greatest sensitivity for that muscle (Figure 3a-b). For this analysis, we did not use a log-ratio, because EMG activity was not scaled similarly in R2 and R3 epochs (Figure 3d-e), and that difference would lead to a mismatched ratio normalisation across epoch, hindering comparisons. The increase in gains with reward was indeed larger in the R3 epoch than in the R2 epoch, indicating that reward increased feedback gains more in the R3 epoch than it did in the R2 epoch (W=112, r=0.82, p=0.023; Figure 3g). Finally, ROC analysis showed that LLR latencies were similar in the rewarded condition compared to the non-rewarded condition (W=73, r=0.54, p=0.48, Figure 3h-i).

In summary, while the prospect of reward did not alter the SLR, it led to increases in feedback gains as early as the LLR, that is, about 50 ms post-perturbation, which is much earlier than the increase in latencies with reward reported in previous work (Carroll et al., 2019; De Comité et al., 2021). This increase in LLR gains with reward was stronger in the R3 epoch than in the R2 epoch.

### 2.3. Latencies for selecting a target were reduced with reward

In addition to the SLR and LLR, slower feedback responses also exist that control for higher level aspects of movement, such as selecting a target based on external cues (Figure 1). We tested the effect of reward on this feedback response in a “Target Selection” task. In that task, participants were instructed to select a target based on the direction of a mechanical perturbation (Figure 4a). Specifically, using the same layout as the previous task measuring the SLR and LLR, half of the trials (112/224) contained two targets, and participants were instructed to reach to the target opposite to the perturbation direction following perturbation onset (Figure 4a, and purple trajectories in Figure 4b). In the other half of trials, only one target was displayed, and participants were instructed to reach to that target following perturbation onset, bypassing the need for any “selection” process (orange trajectories). For both one- and two-target conditions, outward and inward perturbations occurred equally to make the perturbation direction unpredictable.

**Figure 4:**
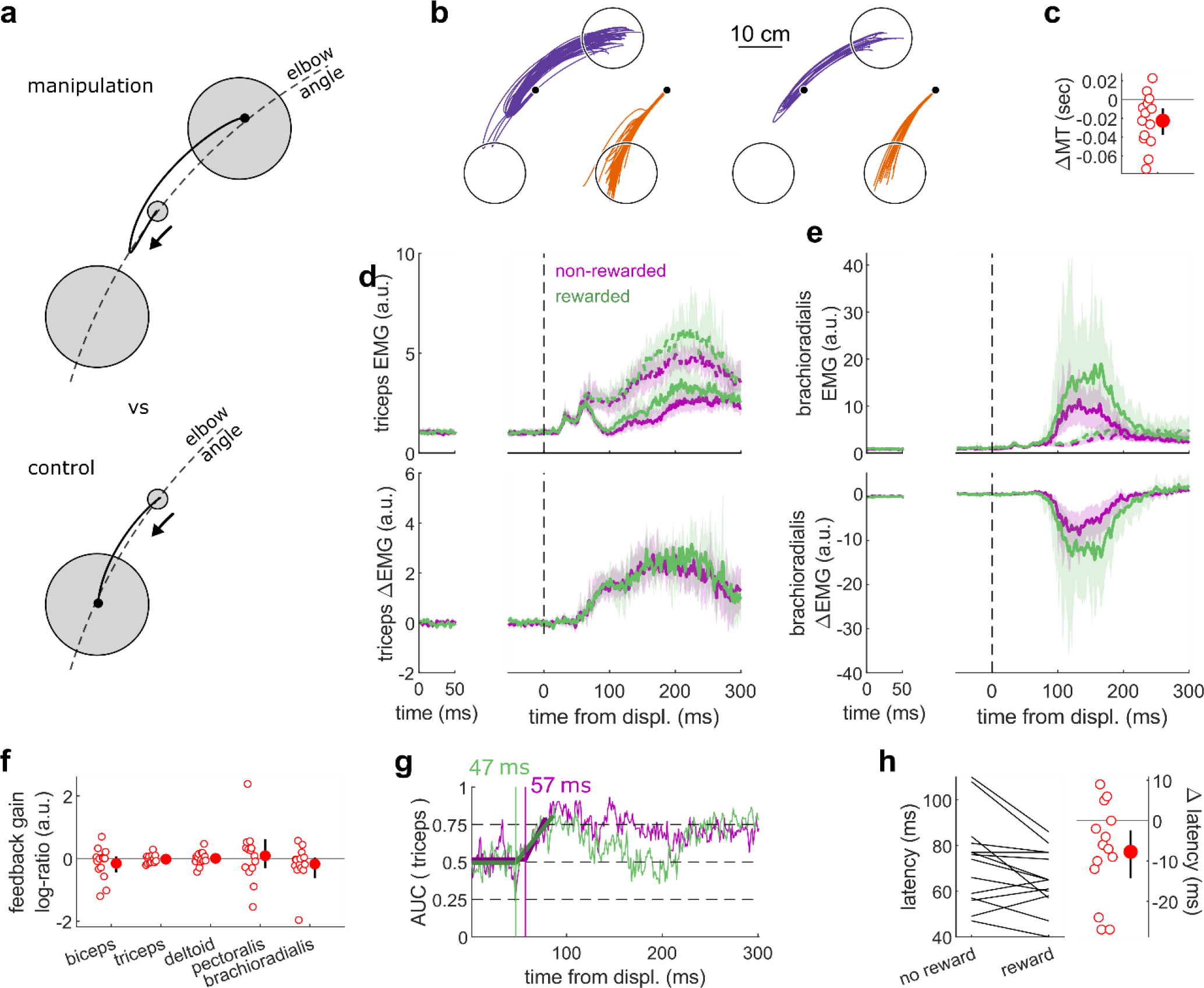
Results for the Target Selection task. **(a)** Contrast used to observe the feedback response to a Target Selection. Background loads are not drawn for clarity. **(b)** Trajectories for one participant (left) and average trajectory for all participants (right, N=14). **(c)** Difference in mean movement time between rewarded and non-rewarded trials. A negative value indicates a smaller movement time for rewarded trials. **(d)** Average triceps EMG signal across participants, with the dashed and solid lines representing manipulation and control conditions, respectively, as indicated in (a); bottom panels: difference between the manipulation and control condition. The left panels show EMG at trial baseline (see methods). Shaded areas indicate 95% Cis. **(e)** Same as (d) but for the brachioradialis muscle. **(f)** Feedback gains following Target Selection response onset. **(g)** Example area under curve (AUC) to obtain response latency for one participant. Thick lines indicate line-of-best-fit for a two-step regression (see methods). **(h)** Response latencies. In all panels with a red filled dot and black error bars, the filled dot indicates the group mean and error bars indicate 95% CIs.

Similar to the previous experiments monitoring the SLR and LLR, participants were rewarded for shorter movement times. We computed for each participant the median MT of trials corresponding to the conditions of interest (Figure 4a) for rewarded and non-rewarded trials and compared them using a Wilcoxon rank-sum test. Performance improved in the rewarding condition (W=92, r=0.88, p=0.011, Figure 4c), and this was not due to an increase in feedback gains (biceps: W=41, r=0.39, p=0.5; triceps: W=68, r=0.65, p=0.36; deltoid: W=54, r=0.51, p=0.95; pectoralis: W=59, r=0.56, p=0.71; brachioradialis: W=70, r=0.67, p=0.3; Figure 4f) but to a shortening of response latencies (W=76, r=0.72, p=0.031, Figure 4g-h).

### 2.4. Reaction times improved with reward

Reaction times have been measured in many different settings that include rewarding feedback (Douglas and Parry, 1983; Steverson et al., 2019; Stillings et al., 1968). The consensus is that reaction times are reduced when reward is available. However, previous work always considered reaction times triggered by non-proprioceptive cues, such as auditory (Douglas and Parry, 1983) or visual cues (Stillings et al., 1968). Here, we assessed participants’ reaction times triggered by a proprioceptive cue, which for arm movement tasks produce faster response latencies than visual cues (Pruszynski et al., 2008).

Participants held their arm so that the tip of the index finger was positioned at a starting location and the arm was stabilized against background loads that pre-loaded the forearm and upper arm extensor muscles (Figure 5a). A go cue was provided in the form of a small flexion perturbation at the shoulder, that led to less than 1 degree of shoulder or elbow rotation (Pruszynski et al., 2008). Participants were instructed to perform a fast elbow extension toward a target (10 cm radius) when they detected the go cue. Median movement times were greatly reduced in the rewarded condition compared to the non-rewarded condition (W=153, r=1, p=2.9e-4, Figure 5c), again indicating that the task successfully increased participants’ motivation. Reaction times were defined as when the (processed) triceps EMG signal rose 5 standard deviations above baseline level (Pruszynski et al., 2008) for 5 ms in a row (Figure 5d). In line with the literature on reaction times triggered by other sensory modalities, proprioception-triggered reaction times were also reduced under reward, reducing on average by 13.7 ms, from 174.8 to 161.1 ms (W=107, r=0.7, p=0.043, Figure 5f). Feedback gains also increased significantly for all recorded muscles (biceps: W=144, r=0.94, p=1.4e-3; triceps: W=150, r=0.98, p=5.0e-4; deltoid: W=150, r=0.98, p=5.0e-4; pectoralis: W=135, r=0.88, p=5.6e-3; brachioradialis: W=138, r=0.9, p=3.6e-3; Figure 5e).

**Figure 5:**
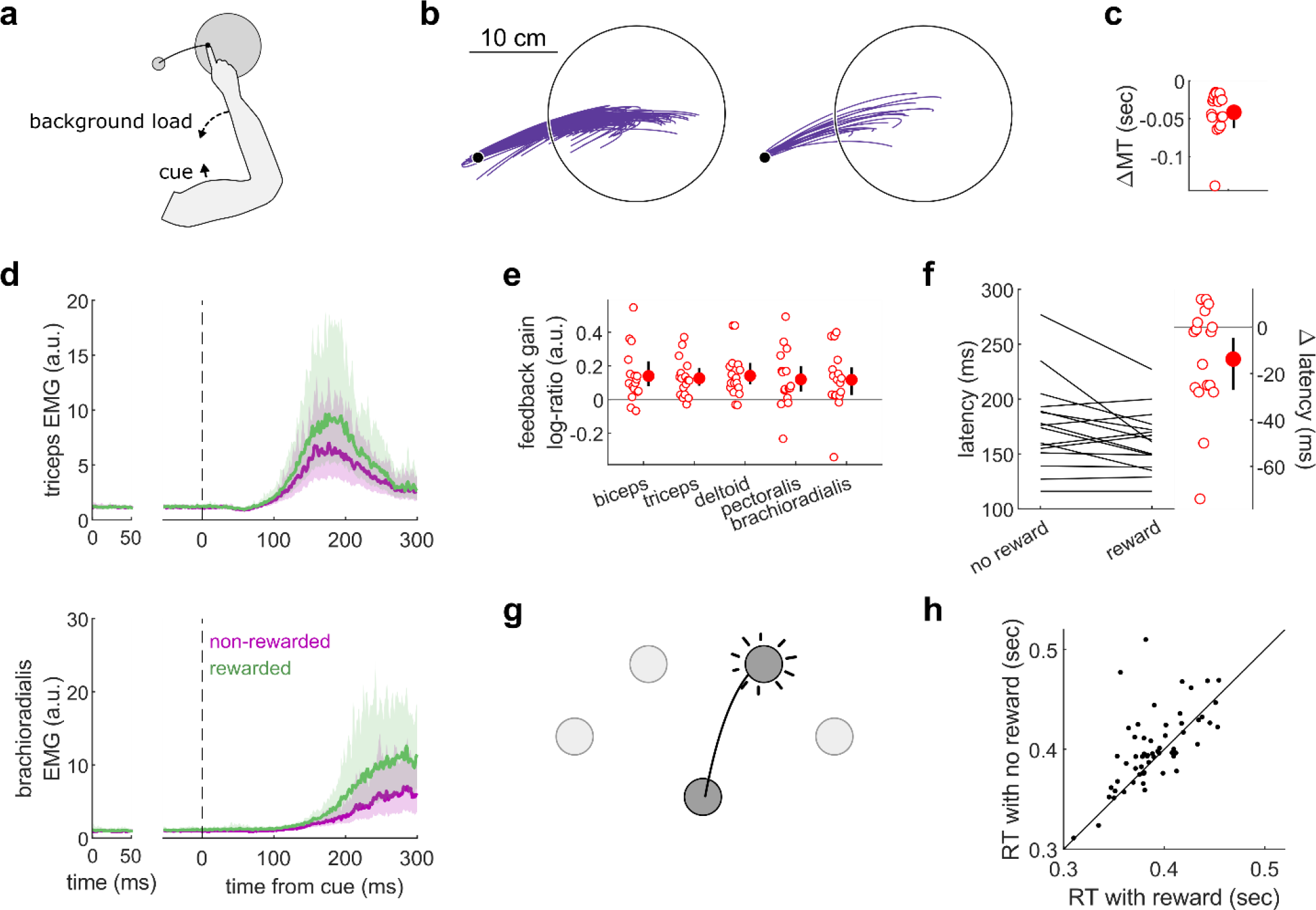
Results for the Reaction Time tasks. **(a)** Schematic of task design for Proprioception-cued Reaction Times. Participants were informed to initiate an elbow extension by a small mechanical perturbation at the shoulder (solid black arrow). Background loads pre-loaded the elbow extensor muscles (dashed black arrow). **(b)** Trajectories for one participant (left) and average trajectory for all participants (right, N=17). **(c)** Difference in mean movement time between rewarded and non-rewarded trials. A negative value indicates a smaller movement time for rewarded trials. **(d)** Top panels: average triceps EMG signal across participants. The left panels show EMG at trial baseline (see methods). Shaded areas indicate 95% CIs. Bottom panels: same as top panels but for the brachioradialis. **(e)** Feedback gains following the feedback response onset. **(f)** Response latencies. In all panels with a red filled dot and black error bars, the filled dot indicates the group mean and error bars indicate 95% CIs. **(g)** Schematic of task design for choice reaction times. **(h)** Median reaction times for each participant (N=60) in the choice reaction time task in the rewarded and non-rewarded conditions, plotted against unity.

Finally, we assessed reaction times in a choice reaction time task by re-analysing a dataset available online (Codol et al., 2020a). In this dataset, participants (N=60) reached to one of four targets displayed in front of them in an arc centred on the starting position (Figure 5g). Participants could obtain monetary reward for initiating their movements quicker once the target appeared (reaction times) and for reaching faster to the target (movement times). In line with the current study, reaction times were shorter in the rewarded than in non-rewarded condition, from 400.8 to 390.2 ms on average (W=1241, r=0.67, p=0.016, Figure 5h). Of note, because EMG recordings were not available for the online dataset, only kinematic data were available, which explains the slower absolute reaction times than reported in other studies (Haith et al., 2015; Summerside et al., 2018).

### 2.5. Online visual control of limb position was unaltered by reward

Next, we assessed feedback response relying on visual cues rather than proprioceptive cues. In a new task, a visual jump of the cursor indicating hand position occurred halfway through the movement, that is, when the shoulder angle was at 45 degrees like the Target Selection task (Figure 6a). To improve tuning of the triceps EMG signal to the feedback response, the reach and jump were specified in limb joint angle space, with the reach corresponding to a shoulder flexion rotation, and the cursor jump corresponding either to an elbow flexion or extension rotation (Figure 6a-b). A third of trials contained no jumps. Like the experiments probing feedback responses relying on proprioception, participants were rewarded for shorter movement times.

**Figure 6:**
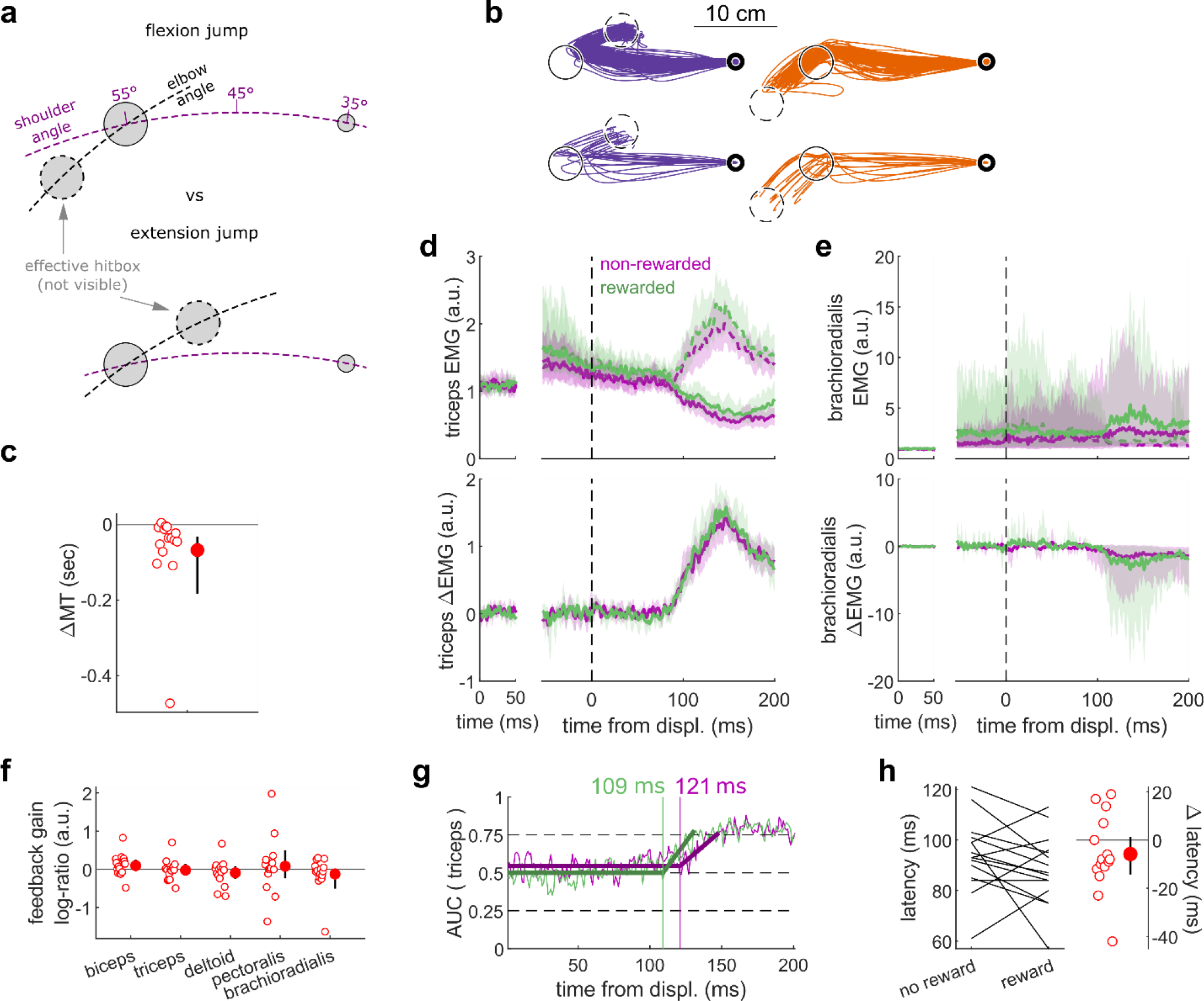
results for the Cursor Jump task. (a) Contrast used to observe the feedback response to a cursor jump. (b) Trajectories for one participant (top) and average trajectory for all participants (bottom, N=15). The dashed circles correspond to where the actual target “hit box” is to successfully compensate for the cursor jump. (c) Difference in mean movement time between rewarded and non-rewarded trials. A negative value indicates a smaller movement time for rewarded trials. (d) Average triceps EMG signal across participants, with the dashed and solid lines representing a flexion jump and an extension jump, respectively; bottom panels: difference between the flexion and extension condition. The left panels show EMG at trial baseline (see methods). Shaded areas indicate 95% CIs. (e) Same as (d) but for the brachioradialis. (f) Feedback gains following the feedback response onset. (g) Example area under curve (AUC) to obtain response latency for one participant. Thick lines indicate line-of-best-fit for a two-step regression (see methods). (h) Response latencies. In all panels with a red filled dot and black error bars, the filled dot indicates the group mean and error bars indicate 95% CIs.

Again, behavioural performance improved with reward, as measured by the difference in each participant’s median movement time in the rewarded versus non-rewarded trials in the conditions of interest (Figure 6a; W=108, r=0.90, p=0.0043, Figure 6c), indicating that the rewarding context successfully increased participants’ motivation. However, this improvement in performance was not driven by an increase in feedback gains (biceps: W=85, r=0.71, p=0.17; triceps: W=70, r=0.58, p=0.6; deltoid: W=76, r=0.63, p=0.39; pectoralis: W=73, r=0.61, p=0.49; brachioradialis: W=74, r=0.62, p=0.45; Figure 6f) nor a latency shortening (W=71, r=0.59, p=0.26, Figure 6g-h).

### 2.6. Feedback gains increased to respond to a visual Target Jump

Finally, we assessed the feedback response arising from a visual shift in goal position using a target jump paradigm. The task design was identical to that of the Cursor Jump task (Figure 6a), except that the target, rather than the cursor, visually jumped in the elbow angle dimension (Figure 7a-b). Performance improved in the rewarding condition as well (W=103, r=0.98, p=3.7e-4, Figure 7c). Unlike for cursor jumps, feedback gains in the Target Jump task increased in the rewarding context for the triceps, pectoralis, and brachioradialis muscles (biceps: W=82, r=0.78, p=0.068; triceps: W=94, r=0.9, p=6.7e-3; deltoid: W=74, r=0.7, p=0.19; pectoralis: W=105, r=1, p=1.2e-4; brachioradialis: W=94, r=0.9, p=6.7e-3; Figure 7f). However, the response latencies remained similar between rewarded and non-rewarded conditions (W=67, r=0.64, p=0.39, Figure 7g-h).

**Figure 7:**
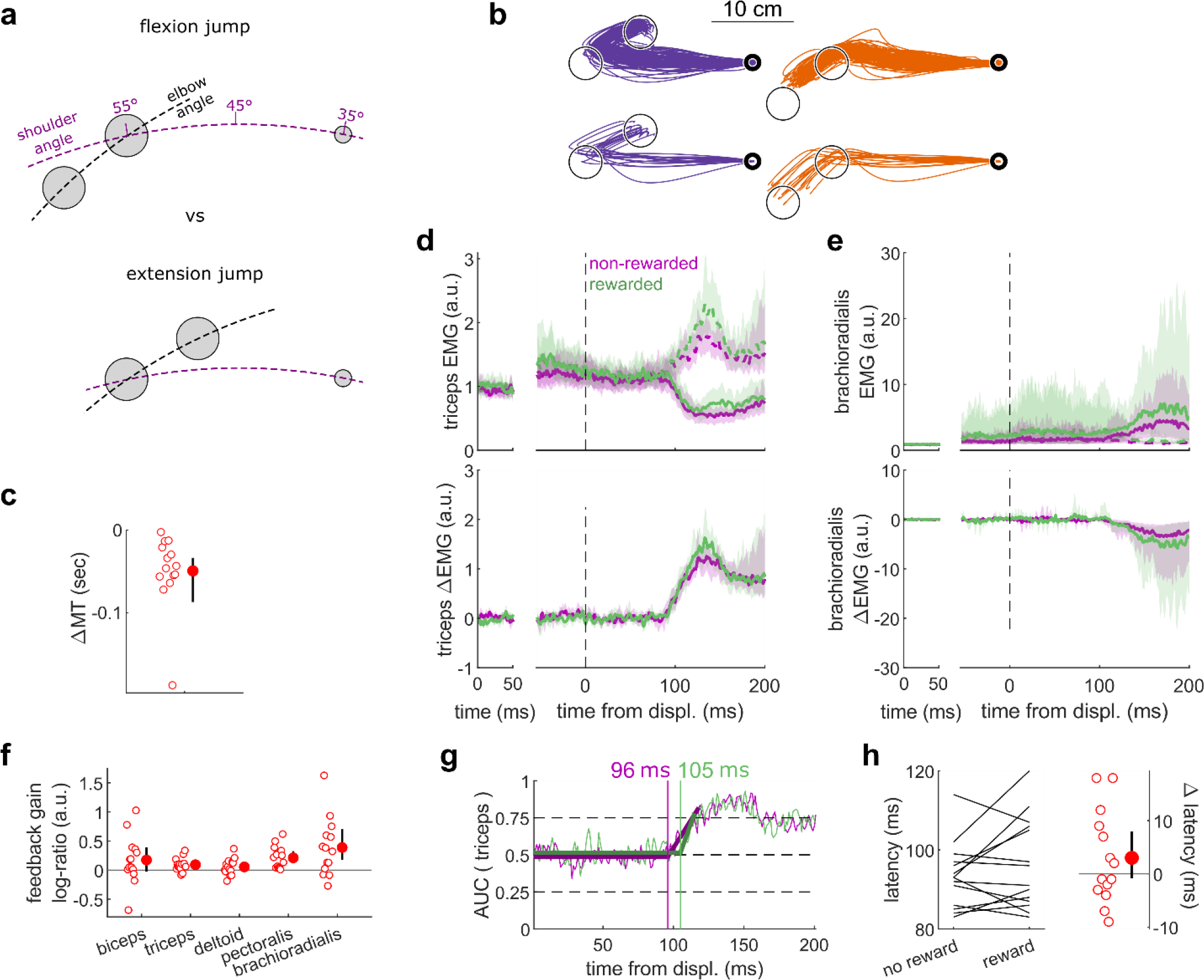
Results for the Target Jump task. **(a)** Contrast used to observe the feedback response to a target jump. **(b)** Trajectories for one participant (top) and average trajectory for all participants (bottom, N=14). **(c)** Difference in mean movement time between rewarded and non-rewarded trials. A negative value indicates a smaller movement time for rewarded trials. **(d)** Average triceps EMG signal across participants, with the dashed and solid lines representing an extension jump and a flexion jump, respectively, as indicated in (a); bottom panels: difference between the extension and flexion conditions. The left panels show EMG at trial baseline (see methods). Shaded areas indicate 95% CIs. **(e)** Same as (d) but for the brachioradialis. **(f)** Feedback gains following the feedback response onset. **(g)** Example area under curve (AUC) to obtain response latency for one participant. Thick lines indicate line-of-best-fit for a two-step regression (see methods). **(h)** Response latencies. In all panels with a red filled dot and black error bars, the filled dot indicates the group mean and error bars indicate 95% CIs.

## 3. Discussion

In this study we tested whether reward affected seven different kinds of sensorimotor feedback responses through five experiments and re-analysing results from an online dataset (central column in Table 1). As expected, results indicate a heterogeneous sensitivity, both in terms of which feedback responses and which characteristics of the response were modulated by reward (Figure 8). The earliest effect was observed during the R2 epoch of the LLR, that is, about 50 ms post-perturbation. This effect was constrained to the gain of the feedback response and did not extend to its latency. Following this, slower feedback responses in the proprioceptive domain were all affected by the prospect of reward (Figure 8). In the visual domain, the Target Jump task, and all slower feedback responses were affected as well. The fastest feedback responses for the proprioceptive and visual domain showed no modulation by reward, as shown by the SLR measurements and the visuomotor responses following cursor jumps, respectively.

**Table 1:**
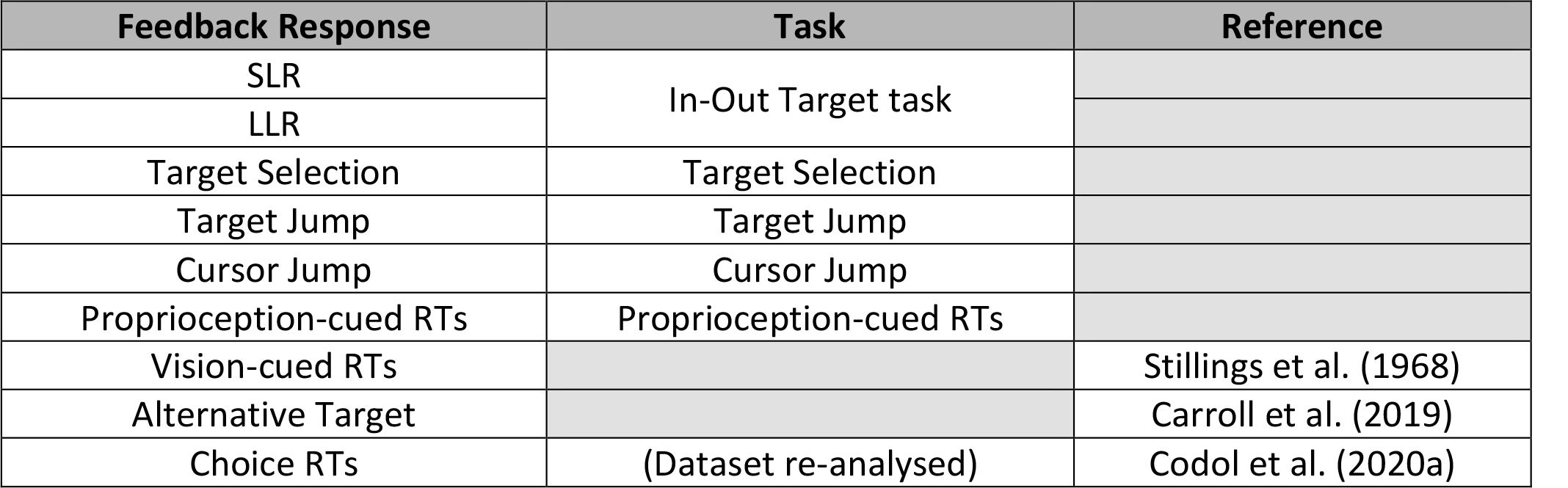
Task to feedback response mapping. This table indicates the correspondence between tasks and published work used, and the feedback response assessed in the present study.

**Figure 8:**
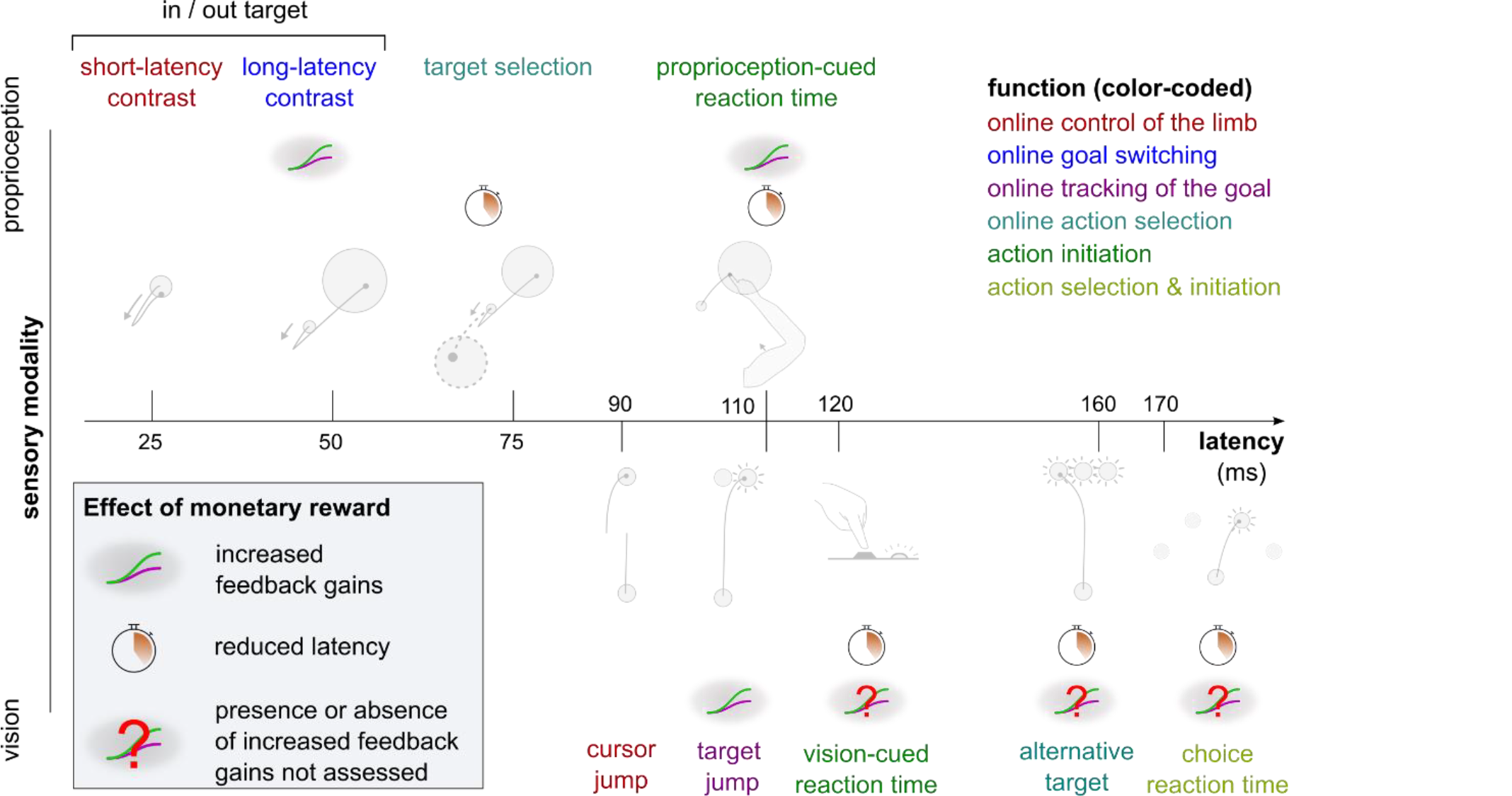
Overview of expected reward impact on sensorimotor feedback responses. Reward can impact a feedback loop response by increasing feedback gains or reducing latency. The colour code indicates function and is identical to the one in Figure 1. Results for the vision-cued reaction time and alternative target tasks are drawn from Carroll et al. (2019) and Stillings et al. (1968), respectively.

### 3.1. Shortening of response latencies may be constrained by transmission delays

The fastest feedback loops showed no reduction in latencies with reward, unlike the slower feedback loops, when adjusted for sensory modality (visual feedback loops tend to be slower than proprioceptive loops). The reproduction of this pattern both in vision and proprioception hints at a mechanism that occurs across sensory domains. Likely, the usual latencies of the fastest feedback loops are constrained by transmission delays. For instance, electrophysiological recordings in monkeys show that proprioceptive information for the LLR takes about 20 ms to reach the primary sensory cortex, and a response travelling back downward would take an additional 20 ms, explaining most of the ∼50 ms latencies generally observed for this feedback loop (Cheney and Fetz, 1984; Omrani et al., 2016; Pruszynski et al., 2011a). Consequently, the LLR has little room for latency improvements beyond transmission delays. This is well illustrated in the Proprioception-cued Reaction Time task, which holds similarities with the task used to quantify the LLR but with a smaller mechanical perturbation. Despite this similarity, latencies were reduced in the Proprioception-cued Reaction Time task, possibly because the physiological lower limit of transmission delays is much below typical reaction times.

As we move toward slower feedback loops, additional information processing steps take place centrally (Carroll et al., 2019; Nashed et al., 2014), which contributes to overall latencies. For instance, the Target Selection and Reaction Time tasks require accumulation of sensorimotor and/or cognitive evidence for triggering movement initiation or selecting an action, respectively. The time course of these processes can vary depending on urgency, utility, and value of information (Fernandez-Ruiz et al., 2011; Reddi and Carpenter, 2000; Steverson et al., 2019; Thorpe and Fabre-Thorpe, 2001; Wong et al., 2017). Therefore, we propose that a rewarding context leads to a reduction in latencies only for the feedback loops relying on online accumulation of sensorimotor and cognitive information, which benefit from that rewarding context. Conversely, typical latencies of the faster feedback loops cannot be shortened because transmission delays cannot be reduced below certain physiological values, regardless of the presence or absence of reward.

### 3.2. Increase in feedback gains through anticipatory pre-modulation

Unlike transmission delays, feedback gains can be adjusted through anticipatory pre-modulation, that is before the movement occurs instead of during the movement (de Graaf et al., 2009; Selen et al., 2012). For instance, the LLR response can vary due to anticipatory cognitive processes such as probabilistic information (Beckley et al., 1991) and verbal instructions (Hammond, 1956). We propose that this capacity to pre-modulate feedback gains may enable the distinct pattern of reward driven improvements compared to latency shortenings (Figure 8).

A corollary to that proposal is that feedback loops that do not show increased feedback gains with reward would also more generally not be susceptible to pre-modulation like we observe in the LLR (Beckley et al., 1991; de Graaf et al., 2009; Takei et al., 2021; Zonnino et al., 2021). A task in our study that is suited to test this possibility is the Cursor Jump task, because it does not show feedback gain modulation, while the Target Jump task, which has a very similar design, does. Therefore, one could consider a version of these tasks in which participants are primed before the trial of the probability of a jump in either direction (Beckley et al., 1991; Selen et al., 2012). In the Target Jump task, the above proposal predicts this should pre-modulate the gain of the feedback response according to the probability value. In the Cursor Jump task it should not.

Capacity for pre-modulation and transmission delays are essentially distinct characteristics. Therefore, an important consequence of our proposal above is that there should be no central coordination between latency reduction and increased feedback gains. For instance, we should not expect feedback gains to increase only when latencies cannot be reduced, that is, as a compensation mechanism. Feedback gains increasing before latencies are reduced may simply occur because it is easier, when possible, to have faster responses via pre-modulation, because it is done before the perturbation occurrence in the first place.

### 3.3. A behavioural classification based on sensory domain and response latency

Several categorization schemes have been used in the literature to sort the large variety of feedback loops involved in feedback control: by typical response latency, by function, or by sensory modality (Figure 1). From our results, categorizing feedback loops by typical latency range appears to be sensible, at least in the context of our observations, and potentially as a general principle. Unsurprisingly, categorization by sensory modality is relevant as well, not only because latency range is impacted by sensory modality, but also because the pathways, neural regions, and mechanisms involved are fundamentally distinct across modalities. This leaves categorization by function (color-coded legend box in Figure 1) outstanding, as it does not match any observed pattern here (Figure 8). This is surprising, as categorization by function is a behavioural classification, and so one may expect it to yield greater explanatory power to interpret results of a behavioural, descriptive set of experiments like we report here. Therefore, while it may have value at a higher-order level of interpretation, we argue that a categorization of feedback loops based on function may not be the most appropriate means of characterizing feedback control loops overall. In contrast, categorization based on neural pathways, neural regions involved, and sensory modality may result in more insightful interpretations.

### 3.4. Implications for Optimal Feedback Control

The Optimal Feedback Control (OFC) framework is a commonly used theoretical framework for explaining how movement is controlled centrally and producing behavioural predictions based on its core principles (Miyashita, 2016; Scott, 2012). One such principle is that feedback gains may be tuned according to a cost function to produce optimal control (Todorov, 2004). The results we report here generally agree with this principle, because we do observe modulation of feedback gains as we manipulate expectation of reward (and so expected cost of movement). However, to our knowledge, previous OFC implementations do not distinguish between individual feedback loops in that regard, that is, it is assumed that any feedback loop can adjust their gains to optimize the cost function (Scott, 2012; Shadmehr and Krakauer, 2008; Todorov, 2005). Our results suggest that not all feedback loops respond in this way, and that the OFC framework’s explanatory power may benefit from distinguishing between the loops that adjust their gains with reward, and those which do not.

### 3.5. General considerations

While SLR circuitry is contained within the spinal cord, it receives supraspinal modulation and displays functional sensitivity to higher-order task goals (Crone and Nielsen, 1994; Nielsen and Kagamihara, 1992, 1993; Weiler et al., 2019). Spinal reflex responses can also be modulated over weeks in the context of motor learning using operant conditioning (Wolpaw, 1987; Wolpaw and Herchenroder, 1990). Therefore, while unlikely, central modulation of SLR circuitry for rewarding outcomes during motor control could not be *a priori* ruled out. However, we observed no such modulation in the present experiments.

A feedback loop that we did not assess is the cortico-cerebellar feedback loop (Becker and Person, 2019; Chen-Harris et al., 2008; Manohar et al., 2019). This loop contributes to saccadic eye movements (Chen-Harris et al., 2008), which show performance improvements under reward as well (Manohar et al., 2015, 2019). Electrophysiological evidence in mice (Becker and Person, 2019) and non-invasive manipulation in humans (Miall et al., 2007) suggest this loop also contributes to reaching movement, but behavioural assessment remains challenging.

### 3.6. Conclusion

Our results combined with previous work (Carroll et al., 2019; Codol et al., 2020a; Douglas and Parry, 1983; Stillings et al., 1968) show that sensitivity to reward is not uniform across all feedback loops involved in motor control. Based on our observations, we propose that (1) reduction of latencies with reward is mainly dictated by neural transmission delays and the involvement (or lack) of central processes in the loop considered, and (2) increase of feedback gains with reward may be the result of central pre-modulation. We also argue against a “top-down” classification of feedback loops based on function, and in favour of a “bottom-up” classification based on neural pathways and regions involved. Finally, we propose potential refinements to apply to the OFC framework based on our results.

Together with previous work showing reduction in peripheral noise with reward (Codol et al., 2020a; Manohar et al., 2019), the results presented here enable us to further complete the picture on how rewarding information triggers improvements in motor performance at the behavioural level. Outstanding questions remain on how reward leads to improvements in motor control, such as whether noise reduction may also occur centrally (Goard and Dan, 2009; Manohar et al., 2015; Pinto et al., 2013), or whether the cortico-cerebellar feedback loop is also involved in reward-driven improvements (Becker and Person, 2019; Codol et al., 2020a; Miall et al., 2007). Beyond motor control, it remains to be tested whether the motor control improvements we observe could be assimilated through motor learning to systematically enhance athletic coaching (Hamel et al., 2019) and rehabilitation procedures (Goodman et al., 2014; Quattrocchi et al., 2017).

## 4. Methods

### 4.1. Dataset and analysis code availability

All behavioural data and analysis code will be made freely available online on the Open Science Framework website at https://osf.io/7t8yj/ upon publication.

### 4.2. Participants

16, 15, 14, 14, and 17 participants took part in the In-Out target, Cursor Jump, Target Jump, Target Selection, and Proprioception-cued Reaction Time task, respectively, and were remunerated CA$12 or 1 research credit per hour, plus performance-based remuneration. Participants made on average 2.14, 2.83, 2.39, 3.80 and 3.19 Canadian cents per rewarded trial on the In-Out Target, Cursor Jump, Target Jump, Target Selection, and Proprioception-cued Reaction Time task, respectively, and earned on average in total $7.19, $4.24, $3.59, $4.26, and $1.72 from performance, respectively. All participants signed a consent form prior to the experimental session. Recruitment and data collection were done in accordance with the requirements of the research ethics board at Western University.

### 4.3. Apparatus

A BKIN Technologies (Kingston, ON) exoskeleton KINARM robot was used for all the tasks presented here. In all cases the participant was seated in front of a horizontally placed mirror that blocked vision of the participant’s arm and reflected a screen above so that visual stimuli appeared in the same plane as the arm. Electromyographic activity of brachioradialis, triceps lateralis, pectoralis major, posterior deltoid, and biceps brachii was recorded using wired surface electrodes (Bagnoli, Delsys, Natick, MA). EMG and kinematic data were recorded at 1000 Hz.

The participant’s arm was rested on a boom that supported the limb against gravity and allowed for movement in a horizontal plane intersecting the centre of the participant’s shoulder joint. Pilot tests using an accelerometer fixed on the distal KINARM boom showed that logged perturbation timestamps corresponding to the onset of commanded robot torque preceded the acceleration of the distal end of the robot linkage by 4 ms. Perturbation timestamps were adjusted accordingly for the analysis of experimental data. For the visual feedback tasks (Cursor and Target Jumps), perturbation onsets were determined using a photodiode attached to the display screen (see *Target Jump and Cursor Jump Tasks* description in “Experimental design” section below for details).

### 4.4. Experimental Design

#### 4.4.1. General Points

Background loads were used to pre-load extensor muscles to improve measurement sensitivity. In all tasks using mechanical perturbations, perturbation magnitudes were added to background loads. For instance, if a background torque load of −2 Nm was applied and a −4 Nm perturbation was specified, then during the perturbation the robot produced a −6 Nm torque.

In all tasks, the start position and the target(s) were the same colour, which was either pink or cyan blue depending on whether the trial was rewarded or non-rewarded. Target colour assignment to reward conditions was counterbalanced across participants.

Target sizes are in cm rather than in degrees because the degree measurements used here are with respect to joint angles. Therefore, a target with same angular size would result in different metric sizes for individuals with longer upper arm or forearm, due to angular projection (Figure 2a).

#### 4.4.2. In-Out Target Task

The location of the tip of the participant’s right index finger was indicated by a 3 mm radius white cursor. At the beginning of each trial, a 3 mm radius start position appeared, along with a reward sign below the target showing “000” or “$$$” to indicate a non-rewarded or a rewarded trial, respectively. The start position was located so that the participant’s external shoulder angle was 45 degrees relative to the left-right axis (parallel to their torso), and the external elbow angle was 90 degrees relative to the upper arm (Figure 1). When participants moved the cursor inside the start position the cursor disappeared. It reappeared if the participant exited the start position before the perturbation onset. After the cursor remained inside the start position for 150-200 ms, a background torque (+2 Nm) ramped up linearly in 500 ms at the shoulder and elbow to activate the extensor muscles. Then, following another 150-200 ms delay a 10 cm radius target appeared either at +20 or −20 degrees from the start position (rotated about the elbow joint). Following target appearance and after a 600 ms delay during which we assessed baseline EMG activity for that trial, the robot applied a ±2 Nm perturbation torque at the elbow and shoulder joints (Figure 2a-c). This combination of load on the shoulder and elbow was chosen to create pure elbow motion, as the robot torque applied at the shoulder counteracted the interaction torque arising at the shoulder due to elbow rotation (Maeda et al., 2018, 2020). Because the time interval between the onset of the visual target and the onset of the perturbation was fixed, we tested for anticipatory triceps EMG activity between the manipulation and control groups in a 20 ms window immediately before the perturbation onset. We observed no difference, both for the SLR (no reward: W=83, r=0.61, p=0.43; with reward: W=88, r=0.65, p=0.30) and the LLR (no reward: W=70, r=0.51, p=0.92; with reward: W=82, r=0.60, p=0.46). Following the mechanical perturbation, participants were instructed to move the cursor as fast as possible to the target and stay inside it until the end of the trial. Each trial ended 800 ms after perturbation onset, at which point the target turned dark blue, the reward sign was extinguished, and the final value of the monetary return was displayed in its place. For non-rewarded trials, this was always “0 ¢” and for rewarded trials, this was calculated as the proportion of time spent in the target from the perturbation onset to the trial’s end:

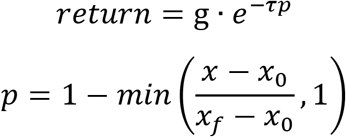

where *x* is the time (ms) spent in the target, *x*_0_ = 500 is the minimum amount of time (ms) to receive a return, *x*_*f*_ = 800 is the total duration (ms) of the trial, *g* = 15 is the maximum return (¢), and *τ* is a free parameter adjusted based on pilot data to reduce the discrepancy between easier and harder conditions. In this study, we used *τ* = 1.428 and *τ* = 2.600 for an inward and outward perturbation with an outward target, respectively, and *τ* = 2.766 and *τ* = 1.351 for an inward and outward perturbation with an inward target, respectively.

The task consisted of 336 trials and was divided into three equal blocks with a free-duration break time between each block. Each block consisted of 112 trials, equally divided between inward and outward perturbation torques, inward and outward target positions, and rewarded and non-rewarded trials. The trial schedule was organised in epochs of 16 trials containing two of each combination of conditions, and the trial order was randomized within each epoch.

For EMG analysis, inward and outward perturbations were used as the manipulation and control conditions, respectively, to observe the SLR on extensor muscles (Figure 2a). To observe the LLR on the extensor muscles, inward perturbations were used when combined with an outward and inward target for the manipulation and control condition, respectively (Figure 3a).

#### 4.4.3. Proprioception-cued Reaction Time Task

The Proprioception-cued Reaction Time task used the same task as the In-Out Target task, with several alterations. First, the background loads were applied to the elbow only, and only outward targets were presented, so that the task consisted of elbow extension movements only. The starting position was located such that the external shoulder angle was 45 degrees relative to the left-right axis, and the external elbow angle was 70 degrees relative to the upper limb. The end target was located at a shoulder and elbow angle of 45 degrees. The perturbation was applied only at the shoulder instead of both shoulder and elbow joints, and the perturbation magnitude was reduced to 0.5 Nm, to ensure that the perturbation led to no significant elbow motion (Pruszynski et al., 2008). Finally, the perturbation time was jittered in a 600-1000 ms window following the target appearance (random uniform distribution). This window was used to measure baseline EMG activity for that trial. Participants were informed to initiate a movement to the target as fast as possible following the shoulder perturbation. Movement times were defined as the time interval from the perturbation occurrence to entering the end target, regardless of velocity. They performed 27 epochs of 2 rewarded and 2 non-rewarded trials randomly interleaved, yielding a total of 108 trials per participant.

Monetary returns were calculated using the following formula:

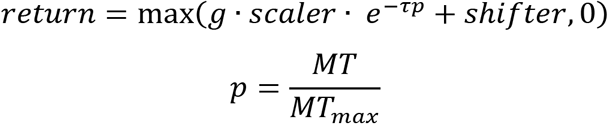

where *MT* is the movement time, *MT*_*max*_ is a normalizing movement time value, *g* = 10 is the maximum amount of money (CAD cents) that may be earned in a trial, and *scaler, shifter, τ* are parameters that allow calibrating the return function to typical psychometric performance in the task considered. Their value was determined using pilot data to ensure large variance across participants based on performance and are provided in Table 2.

**Table 2:** Parameters used to compute the return in each rewarded trial, for each condition in the Proprioception-cued Reaction Time, Cursor Jump, Target Jump, and Target Selection tasks.

| Task | Condition | Scaler | Shifter | $\tau$ | MT<sub>max</sub> [ms] |
| --- | --- | --- | --- | --- | --- |
| Reaction Time | N/A | 1 | 0 | 2.447 | 728 |
| Cursor Jump | Inward | 0.996 | -0.029 | 4.273 | 2781 |
|  | No jump | 0.667 | 0.079 | 5.433 | 2335 |
|  | Outward | 0.996 | -0.041 | 3.958 | 2864 |
| Target Jump | Inward | 0.999 | -0.026 | 4.281 | 2697 |
|  | No jump | 0.683 | -0.040 | 3.893 | 2882 |
|  | Outward | 0.999 | -0.054 | 3.853 | 2690 |
| Target Selection | One target Inward Pert. | 0.676 | -0.034 | 6.236 | 1673 |
|  | One target Outward Pert. | 0.690 | 0.004 | 5.534 | 2241 |
|  | Two targets Inward Pert. | 0.749 | -0.021 | 4.904 | 2373 |
|  | Two targets Outward Pert. | 0.749 | 0.009 | 5.350 | 2208 |

For the choice reaction time task, the methods employed are described in Codol et al. (2020a).

#### 4.4.4. Target Jump and Cursor Jump Tasks

The position of the participants’ right hand index fingertip was indicated by an 8 mm radius white cursor. At the beginning of each trial, an 8 mm radius start position was displayed at a shoulder and elbow angle of 35 degrees and 65 degrees, respectively, and below it a reward sign showing “000” or “$$$”to indicate a non-rewarded or a rewarded trial, respectively. At the same time, a 2 cm radius target appeared at a 55 degrees shoulder angle and same elbow angle as the start position (65 degrees). This yields a reaching movement that consists of a 20 degrees shoulder flexion with the same elbow angle throughout (Figure 7a, 6a). Participants moved the cursor inside the start position, and after 200 ms, +2 Nm shoulder and elbow background torques ramped up linearly in 500 ms. Participants held the cursor inside the start position for 600-700 ms (random uniform distribution), during which baseline EMG activity for that trial was measured. Following this, the end target appeared. Participants were instructed to reach as fast as possible to that end target, and that their movement time would define their monetary return on “$$$” trials but not “000” trials. They were informed that reaction times were not considered into the calculation of the return. Movement duration was defined as the time interval between exiting the start position and when the cursor was inside the target and its tangential velocity dropped below 10 cm/sec. Once these conditions were met, the target turned dark blue, the reward sign was extinguished, and the final monetary return for the trial appeared where the reward sign was located before. In the Target Jump task, during the reach when the cursor crossed past the (invisible) 45 degrees shoulder angle line, the target jumped to ±10 degrees elbow angle from its original position or stayed at its original position (no-jump), with all three possibilities occurring with equal frequency (Figure 7a). In the Cursor Jump task, the cursor position rather than the target jumped to ±10 degrees elbow angle or did not jump.

Monetary return was given using the same equation as for the reaction time task, with *g* = 10. The parameters used for each condition were calibrated using pilot data to ensure similar average returns and variance across conditions within participants. The values used are provided in Table 2.

Both the target and cursor jumps consisted of 312 trials in one block, equally divided between rewarded and non-rewarded trials, and outward jump, inward jump, and no-jump trials in a 2×3 design. The trial schedule was organised in epochs of 12 trials containing two of each combination of conditions, and the trial order was randomized within each epoch.

For EMG analysis, outward target jumps, and inward cursor jumps were used as the manipulation condition because they both led to contraction of the triceps muscle to adjust the movement. Inward target jumps and outward cursor jumps were used as control conditions. No-jump conditions were not used for EMG analysis. All EMG signals were aligned to a photodiode signal recording the appearance of a white 8 mm radius target at the same time as the jump occurrence. The photodiode target was positioned on the screen horizontally 25 cm to the right from the starting position and vertically at the same position as the cursor jumping position (in the cursor jump task) or as the target position (target jump task). This target was covered by the photodiode sensor and was therefore not visible to participants.

#### 4.4.5. Target Selection Task

The position of participants’ right index fingertip was indicated by a white 3 mm radius cursor. At the beginning of each trial, a 3 mm radius start position appeared, and a reward sign showing “000” or “$$$” was displayed to indicate a non-rewarded or a rewarded trial, respectively. The start position was located so that the external shoulder angle was 45 degrees relative to the left-right axis, and the external elbow angle was 90 degrees relative to the upper arm. When participants moved the cursor inside the start position the cursor disappeared. it reappeared if the cursor exited the start position before the target(s) appeared. Once inside the start position, the robot applied +2 Nm background torques which were ramped up linearly in 500 ms at the shoulder and elbow to activate the extensor muscles. Then, following a delay of 400-500 ms, a 7 cm radius target appeared at +30 degrees (inward) from the start position (rotated about the elbow joint). In half of trials, a second, identical target also appeared at −30 degrees (outward) from the start position. A jittered 600-1000 ms window followed target appearance, during which baseline EMG activity for that trial was measured. After this, a +2 Nm or −2 Nm perturbation at the shoulder and elbow joints pushed participants away from the start position. Positive and negative perturbations occurred evenly in one- and two-targets trials, yielding a 2×2 task design. For one-target trials, participants were instructed to reach as fast as possible to the target available once the perturbation was applied. For two-target trials, participants were instructed to reach as fast as possible to the target opposite to the perturbation, *e*.*g*., if the perturbation resulted in an inward push, then the participant should go to the outward target. Therefore, this task design resulted in a co-occurrence of the target selection and divergence of triceps EMG activity compared to one-target trials, enabling us to assess the feedback response that underlies selection of the goal target.

When the (correct) end target was reached, the target(s) turned dark blue, the reward sign was extinguished, and the final monetary return for the trial appeared where the reward sign was located before. For non-rewarded trials, this was always “0 ¢” and for rewarded trials, the return was higher for shorter movement times. Movement times were defined as the time interval from the perturbation occurrence to entering the (correct) end target, regardless of velocity. The monetary return equation was identical to that of the reaction time task, with *g* = 15 and the other parameters as provided in table 2. These parameters were calibrated using pilot data to ensure similar average returns and variance across conditions within participants.

The task consisted of 224 trials and was divided into 2 blocks with a free-duration break time between each block. Each block consisted of 112 trials, equally divided between one- and two-targets trials, inward and outward perturbation trials, and rewarded and non-rewarded trials. The trial schedule was organised in epochs of 16 trials containing two of each combination of conditions, and the trial order was randomized within each epoch.

### 4.5. EMG signal processing

EMG signals were sampled at 1000 Hz, band-pass filtered between 20 Hz and 250 Hz, and then full wave rectified. Before each task, participants’ baseline EMG signal was acquired by asking participants to position their arm such that the cursor remained, motionless, at the start position for 2 seconds (against the background load, if applicable for the task). This was repeated 4 times, after which the task started normally. Following band-pass filtering and full-wave rectification, the EMG signal of each muscle from 250 ms after entering the start position to 250 ms before the end of the 2 seconds window was concatenated across all 4 trials and averaged to obtain a mean baseline EMG scalar. EMG measures during the task were then normalised by each muscle’s baseline scalar. Follow-up analyses (latency, feedback gains, co-contraction) were performed subsequently on the filtered, full wave rectified and normalised EMG traces.

For all experimental designs, trial-by-trial baseline EMG activity was measured in a 50 ms window from 350 to 300 ms before displacement onset, while participants were at rest, background loads were applied, and after targets and reward context were provided. For the Target Jump and Cursor Jump tasks, this was measured in a 50 ms window from 350 to 300 ms before target appearance instead of before displacement onset because movements were self-initiated, and displacements occurred during the movement. However, the same target was displayed in every condition at the start of a given trial in those two experimental paradigms. Note that these trial-by-trial baseline EMG signals are distinct from the 4 baseline trials described above in this section, which were done before the task started and were used for normalization of EMG signals. The trial-by-trial baseline EMG signals were not used for EMG normalization.

### 4.6. Statistical analysis

To determine the time at which EMG signals for different task conditions diverged, we used Receiver operating characteristic (ROC) analysis. We used the same approach as in Weiler et al. (2015), using a 25-75% threshold of area under the curve (AUC) for establishing signal discrimination. The threshold was considered reached if two consecutive samples were greater than the threshold value. Discrimination was done for each participant and each reward condition independently, using all trials available for each contrast without averaging. Once the AUC threshold was crossed, we performed a segmented linear regression on the AUC before it crossed the 25-75% threshold. We minimized the sums-of-squared residuals to find the inflexion point, that is, where the two segments of the segmented linear regression form an angle (see Weiler et al. (2015) and analysis code online for details).

To compute feedback gains, for each feedback response considered we defined a 50 ms window that started at that response’s latency found for each participant independently using ROC analysis. For the SLR contrast only, we constrained that window to the R1 epoch (Pruszynski et al., 2008), which corresponds to a 25 ms window from the response latency. We then calculated the integral of average EMG signals in that window using the trapezoid rule (MATLAB’s built-in *trapz* function), for the control and manipulation condition and each reward value. For instance, for the Target Selection task the control condition is defined as trials with only one target (no switch), while the manipulation condition is defined as trials with an inward perturbation and two targets (switch occurring). We then calculated the absolute difference between those two conditions as a measure of feedback gains. We then calculated the log-ratio of the rewarded to non-rewarded conditions as log(*rewarded gain/non rewarded gain*). We used ratios to ensure that changes in feedback gains are normalized within participants to EMG activity in the non-rewarded condition. The log function was then applied to linearize the ratio values.

For the R2 and R3 epochs of the LLR, the 25 ms window was defined as the 1^st^-to-25^th^ ms and 26^th^-to-50^th^ ms window post-response, respectively. To test for the epoch-reward interaction between R2 and R3, the rewarded gain and non-rewarded gain for each epoch was used directly instead of computing their log-ratio, since absolute EMG values greatly vary between those two epochs and would lead to an imbalanced normalization.

To test for differences between conditions we used Wilcoxon signed-rank tests. For each test, we reported the test statistic *W*, the effect size *r* (Kerby, 2014) and the p-value.

## Contributions

OC conceptualized and designed the research question; OC, MK, CJF, JMG, JAP, and PLG designed the experiments; OC implemented the tasks, OC and MK collected and analysed the datasets; OC, MK, CJF, JMG, JAP, and PLG interpreted the results; OC made the figures and wrote the first draft; OC, MK, CJF, JMG, JAP, and PLG edited and approved the final version of the manuscript.

## Acknowledgements

This work was supported by the Natural Science and Engineering Council of Canada (RGPIN-2018-05458 to PLG) and the Canadian Institutes of Health Research (PJT-156241 to PLG). We thank Jonathan M. Michaels for helpful comments and suggestions.

